# Effects of Three Natural Dietary Compounds on Insect Pests

**DOI:** 10.1101/2025.05.23.655814

**Authors:** Tomas Cabello, Manuel Gamez

## Abstract

Effective and sustainable pest management is a critical challenge in agriculture. This study evaluated the relative efficacy of three natural compounds —neem oil, boric acid, and gallotannin acid— against larvae of three pest species: *Rhynchophorus ferrugineus* (F.) (red palm weevil), *Plodia interpunctella* (Hübner) (Indian meal moth), and *Spodoptera exigua* (Hübner) (beet armyworm). Laboratory bioassays revealed distinct differences in larval survival and mortality across the species. Neem oil demonstrated high effectiveness against *S. exigua* larvae, achieving 100% mortality at all tested dietary concentrations, and moderate effectiveness against *R. ferrugineus*. In contrast, it had no significant effect on *P. interpunctella*. Boric acid was proved to be universally effective, leading to 100% mortality in all three species at the highest concentration. Gallotannin acid had species-specific effects on *S. exigua* but had negligible effects on *R. ferrugineus* and *P. interpunctella*. These findings highlight the potential of natural, low-toxicity compounds as alternatives to synthetic insecticides. Additionally, this study suggests new frontiers for investigating the role of the gut microbiota in mediating the effectiveness of these compounds. While the results provide promising insights into environmentally friendly pest control strategies, further research, including field trials and microbiological assessments, is needed to fully validate their application in agricultural settings. This work contributes to advancing sustainable pest management practices that reduce environmental impact while addressing critical agricultural needs.

## 1. Introduction

The red palm weevil *Rhynchophorus ferrugineus* (F.) (Col.: Curculionidae) is a pest species that affects various palm and coconut trees in more than 50 countries (CAB, 2021). Its biology, ecology, and control have been reviewed extensively (e.g., Martin & Cabello, 2005; FAO, 2020). The Indian meal moth *Plodia interpunctella* (Hübner) (Lep.: Pyralidae) is a species that affects mainly stored plant products in 47 countries (CAB, 2005a). Biology, ecology, and control of the species have been reviewed by Mohandass et al. (2007). The beet armyworm *Spodoptera exigua* (Hübner) (Lep.: Noctuidae) is a pest affecting herbaceous and horticultural crops in 77 countries (CAB, 2005b). Its biology, ecology, and control have been extensively reviewed (e.g., Belda-Suarez, 1994; Xia-Lin et al., 2011; Garay et al., 2015).

The larvae of these species exhibit distinct feeding behaviors and locations, thus impacting control methods. Synthetic organic insecticides are commonly used for chemical control (*e.g*., Hernandez et al., 2003; Campos & Phillips, 2010; Ishtiaq et al. 2021). This has resulted in all three species developing resistance to various active insecticidal ingredients. (e.g., Attia et al., 1979; Ahmed & Freed, 2021; Ishtiaq et al., 2021; Rabelo et al., 2022).

Research on natural plant-derived products for managing these three pest species is limited (e.g., Mantzoukas et al., 2020; Chen et al., 2021; Yan et al., 2021; Moullamri et al., 2025).

Boric acid and borate salts are naturally occurring compounds found in rocks, soil, plants, and water, derived from the element boron (NPIC, 2025). These substances serve as active ingredients in pesticides, effectively targeting insects, spiders, mites, algae, molds, fungi, and weeds (NPIC, 2025). Boric acid is an effective biocide with bactericidal, fungicidal, and virucidal properties (McDonnell, 2017; Celikezen & Sahin, 2023). It is used as an insecticide by adding a food attractant to dry powder, which is then sprinkled into cracks. Insects pick up the powder on their legs and ingest it while cleaning themselves, leading to death by starvation or dehydration within 3‒10 days. The exact action of boric acid on insects is still not fully understood (See et al., 2010; EPA-RED, 2024).

Tannins are complex polyphenols, including hydrolysable tannins, proanthocyanidins, and phlorotannins (Suvanto et al., 2017). They bind and precipitate proteins (Bule et al., 2020). Gallotannin acid, also known as tannic acid, is the most well-known hydrolysable tannin (Singh et al., 2021). Tannins are found naturally in the bark of trees such as sumac, oak, and spruce, and in other plant parts (Khanbabaee & van Ree, 2001). Polyphenols play a role in plant‒insect interactions and defenses, including their structure, regulation, and anti-feeding (antixenosis) and toxicity effects (antibiosis) (Singh et al., 2021). It has been reported that dietary tannins can reduce growth and fecundity in some insect species. However, few studies have identified clear physiological or toxicological impacts; some suggested that tannins might even be positive nutritional factors (Schultz,1989).

Triterpenoids are a wide, biologically interesting group of terpenoids and include a large structural diversity of secondary terpenoids from terrestrial and marine living organisms (Morimoto et al., 1999). Azadirachtin, which is a tetranortriterpenoid, is an active ingredient of neem (*Azadirachta indica* A. Juss.) seed oil. It controls two hundred species of insects, including locusts, gypsy moths, cockroaches, and fall armyworms (Yu, 2008). In insects, this compound interacts with the corpus cardiacum, inhibiting the function of molting hormone. Consequently, it acts as an insect growth regulator by suppressing fecundity, molting, pupation, and adult formation (Ishaaya, 2002). Neem oil has been commercialized as an insecticide and is used globally (Koul, 2023).

The symbiosis between microorganisms and insects, especially the gut microbiota, is crucial for latter’s nutrition, growth, metabolism, homeostasis, and immunity (Yasika & Shivakumar, 2025). However, there are few studies on the effectiveness of natural products, which act as insecticides, with the effects of the gut microbiota on their effectiveness.

This work aimed to determine the relative effectiveness of three low-toxicity natural compounds and to study their effects on three insect pest species, each of which exhibits different feeding behavior.

## 2. MATERIALS & METHODS

The insects (*R. ferrugineus*, *P. interpunctella*, and *S. exigua*) used in these trials were obtained from colonies maintained at the University of Almeria’s Agricultural Entomology Laboratory. The larvae were reared on an optimal artificial semi‒diet composed of agar (20 g), distilled water (880 ml), brewer’s yeast (50 g), wheat germ (50 g), corn meal (50 g), ascorbic acid (4.5 g), coconut fiber (8 g), and a vitamin/amino acid additive (50 ml) (Hidro Rex Vital Aminoacidos ®, S.P. Veterinaria, S.A., Tarragona, Spain). The diet provided 15.0% crude protein (as a percentage of dry weight).

The rearing of *R. ferrugineus* was conducted following the protocol established by Martin & Cabello (2006), whereas that of *P. interpunctella* and *S. exigua* followed the method of Cabello et al. (1984).

The three compounds tested were boric acid (99.5%, Panreac Quimica S.L.U., Barcelona, Spain), gallotannin acid (tannic acid 99.5%, Panreac Quimica S.L.U., Barcelona, Spain), and neem oil (azadirachtin, 3.2% LS, Sipcam, Valencia, Spain). Chemical compounds were added to the semi‒synthetic artificial diet at the specified concentration during preparation.

The trials used a completely randomized design with repeated measures. Each II-instar larva in a 20 ml Coulter vial (20 ml) was an experimental unit. Treatments included five neem oil concentrations (0, 6.3, 12.5, 25.0, 50.0 ppm), six boric acid concentrations (0, 312.5, 625.0, 1,250.0, 2,500.0, 5,000.0 ppm), and seven gallotannin acid concentrations (0, 156.3, 312.5, 625.0, 1,250.0, 2,500.0, 5,000.0 ppm). There were 20 repetitions per treatment (concentration), totaling 100 experimental units for neem oil trial, 120 for boric acid trial, and 140 for gallotannin acid trial, meeting Mead’s resource equation “E” values (Mead, 1988; Mead et al., 2012). Larval survival was tracked at intervals: 3, 6, 9, 12, 15, and 20 days for *R. ferrugineus*, and 3, 6, 9, 12, and 15 days for the other species. Trials were conducted at 25±2 °C, 65±10% R.H. and photo period of 0:24 hours light: dark on *R. ferrugineus* and *P. interpunctella* species, and 16:9 hours on *S. exigua* species.

Larval survival was analyzed by the Kaplan-Meier method with subsequent comparisons between treatments carried out via the long-rank test in each trial. These analyses were performed via IBM SPSS software, version 28. Effective median lethal concentrations (LC_50_) were calculated via probit analysis (Statgraphics statistical analysis software Centurion, version 19).

If there is natural mortality in the controls, adjusted mortality is used according to Abbott’s formula (Abbott, 1925) as follows (Busvine, 1957; Perry et al., 1998):

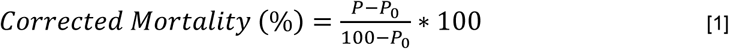

where corrected mortality (%) represents the effectiveness of the tested compound, *P* represents the percentage of mortality of treated insects and *P_0_* represents the percentage of mortality of insects in the untreated control. This value of corrected mortality is accurate if the control mortality is under 20% or is derived from multiple replications (Perry et al., 1998). Piepho et al. (2024) most recently critiqued Abbott’s equation for assuming homogeneity of variance and normality and suggested a generalized linear model (GZLM) to better handle distribution issues. These values were analyzed via GZLM with a Poisson error distribution and logarithm as the link function, considering dietary concentration as a variable factor and days as time-dependent covariate. The analysis was performed using IBM SPSS software (version 28).

## 3. RESULTS

The results for the three compounds analyzed in the three species are shown in the following two sections: the first relating to larval survival and the second, if a response is obtained, measured by the corrected mortalities with respect to the controls (concentrations = 0).

### 3.1. Larval survival

The survival rates of the *R. ferrugineus* larvae were significantly influenced by the dietary concentration of the tested compounds: neem oil (log rank: chi-square = 115.979, df = 4, P < 0.001), boric acid (log rank: chi-square = 321.992, df = 5, P < 0.001), and gallotannin acid (log rank: chi-square = 36.000, df = 6, P < 0.001) (**Fig. 1 a**, **b**, **c**).

**Figure 1.**
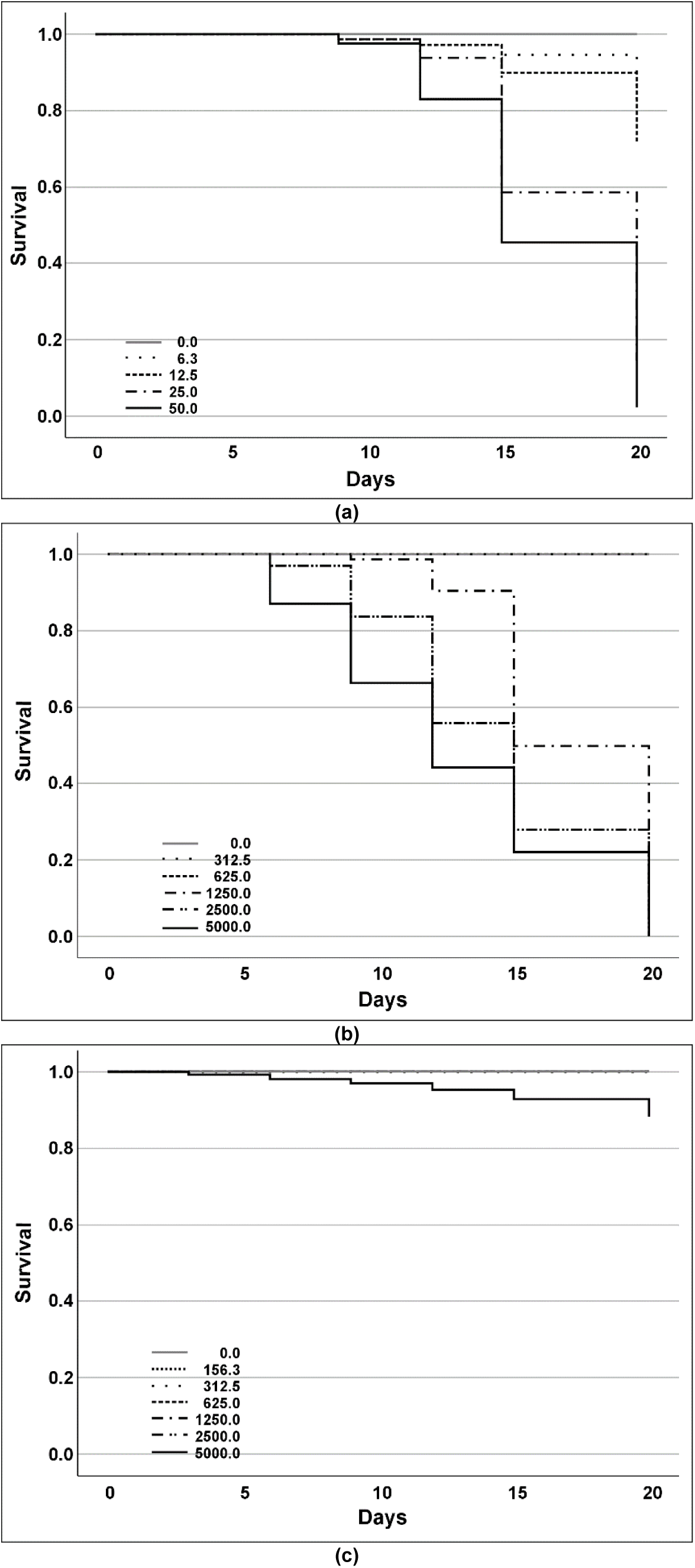
Kaplan–Meier curves and log-rank test for *Rhynchophorus ferrugineus* larval survival when larvae were fed a semi‒synthetic artificial diet containing: (a) neem oil (azadirachtin), (b) boric acid, or (c) gallotannin acid at different concentrations (ppm) under laboratory conditions (25 °C and 0:24 hours light/dark). Control (concentration = 0) (Kaplan–Meier survival analysis, log rank test for overall comparison, P > 0.05).

The effective median lethal concentration (LC_50_) values were 20.6 ppm (lower confidence limit = 16.6 ppm, upper confidence limit = 26.3 ppm, at P = 0.05) at 20 days after treatment (DAT) for neem oil (likelihood ratio tests: chi-square = 64.402, df = 1, P < 0.001) and 1524.1 ppm for boric acid (lower confidence limit = 1338.1 ppm, upper confidence limit = n/a, at P = 0.05) at 12 DAT (likelihood ratio tests: chi-square = 146.172, df = 1, P < 0.001). This could not be estimated for gallotannin acid (likelihood ratio tests: chi-square = 3.626, df = 1, P = 0.057).

Only boric acid significantly affected the larval survival of *P. interpunctella* (log rank: chi-square = 234.331, df = 5, P < 0.001), while neem oil (log rank: chi-square = 0.419, df = 4, P = 0.517) and gallotannin acid (log rank: chi-square = 0.036, df = 6, P = 0.850) did not (**Fig. 2**).

**Figure 2.**
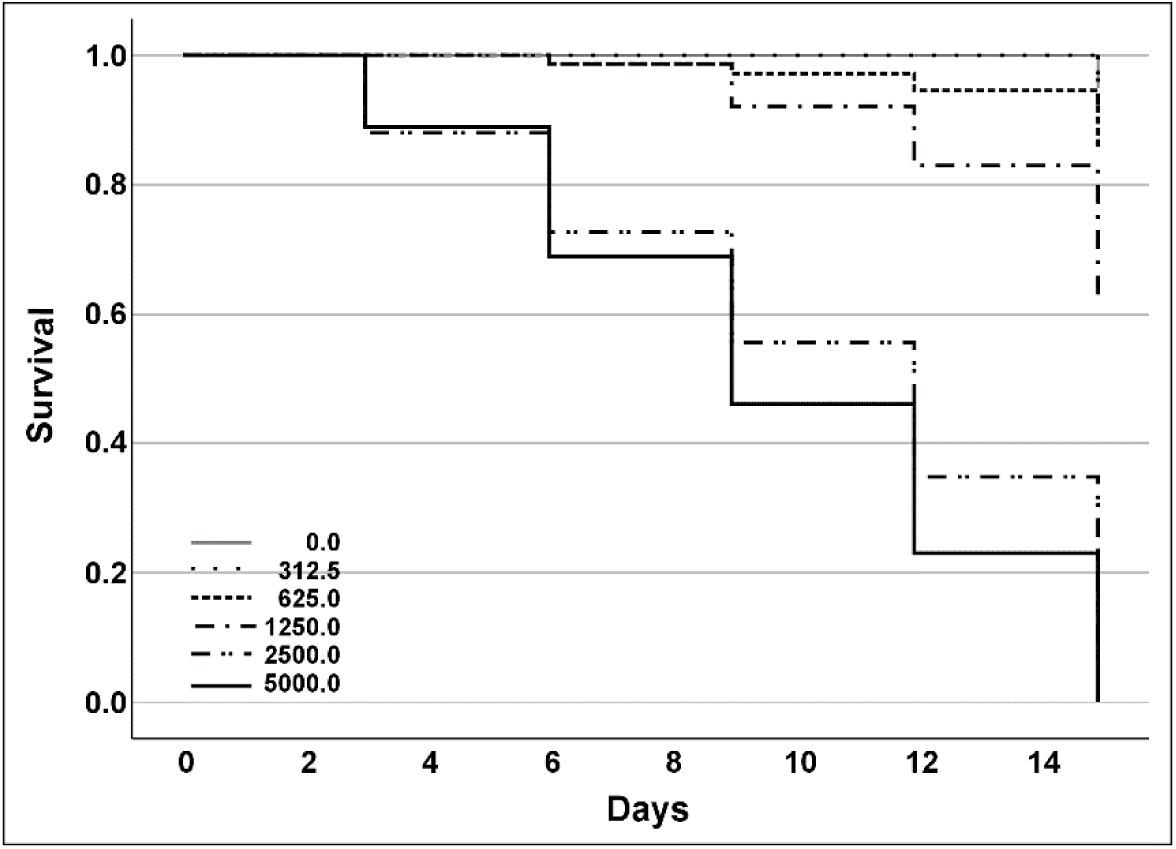
Kaplan–Meier curves and log-rank test for *Plodia interpunctella* larval survival when larvae were fed a semi‒synthetic artificial diet containing boric acid at different concentrations (ppm) under laboratory conditions (25 °C and 0:24 hours Light/Dark). Control (concentration = 0) (Kaplan–Meier survival analysis, log rank test for overall comparison, P > 0.05).

The effective median lethal concentration (LC_50_) value for boric acid was 1760.2 ppm (lower confidence limit = 2403.2 ppm, upper confidence limit = 3767.7 ppm, at P = 0.05) at 9 DAT (Likelihood ratio tests: Chi-square = 89.8753, df = 1, P < 0.001).

The survival rates of the *S. exigua* larvae were significantly influenced by the dietary concentration of the tested compounds: neem oil (log rank: chi-square = 84.144, df = 4, P < 0.001), boric acid (log rank: chi-square = 93.161, df = 5, P < 0.001), and gallotannin acid (log rank: chi-square = 322.096, df = 6, P < 0.001) (**Fig. 3a**, **b**, **c**). The effective median lethal concentration (LC_50_) values for neem oil were 22.7 ppm (lower confidence limit = 18.6 ppm, upper confidence limit = 50.3 ppm, P = 0.05) at 9 DAT (likelihood ratio tests: chi-square = 74.081, df = 1, P < 0.001), 426.1 ppm for boric acid (lower confidence limit = 210.7 ppm, upper confidence limit = 620.5 ppm, P = 0.05) at 9 DAT (likelihood ratio tests: chi-square = 58.631, df = 1, P < 0.001), and for gallotannin acid 1782.4 ppm for gallotannin acid (lower confidence limit = 1473.2 ppm, upper confidence limit = 2237.7 ppm, P = 0.05) at 9 DAT (likelihood ratio tests: chi-square = 96.660 df = 1, P < 0.001).

**Figure 3.**
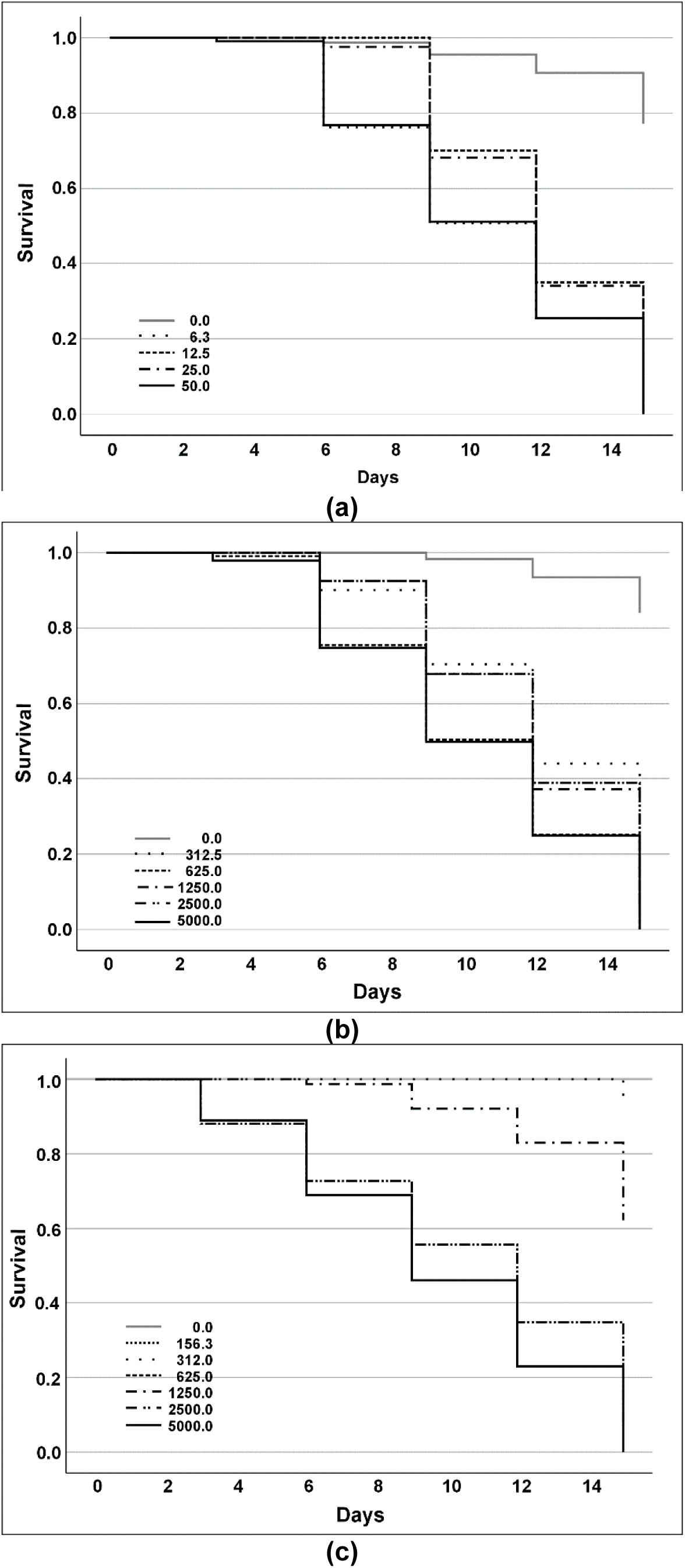
Kaplan–Meier curves and log-rank test for *Spodoptera exigua* larval survival when larvae were fed a semi‒ synthetic artificial diet containing: (a) neem oil (azadirachtin), (b) boric acid, or (c) gallotannin acid at different concentrations (ppm) under laboratory conditions (25 °C and 16:8 hours light/dark). Control (concentration = 0) (Kaplan–Meier survival analysis, log rank test for overall comparison, P > 0.05).

### 3.2. Corrected mortality

**Fig. 4** presents the mortality effectiveness percentages for the *R. ferrugineus* larvae, which were (calculated via Abbott’s formula) [1], when fed with neem oil (**Fig. 4 a**) or boric acid (**Fig. 4 b**).

**Figure 4:**
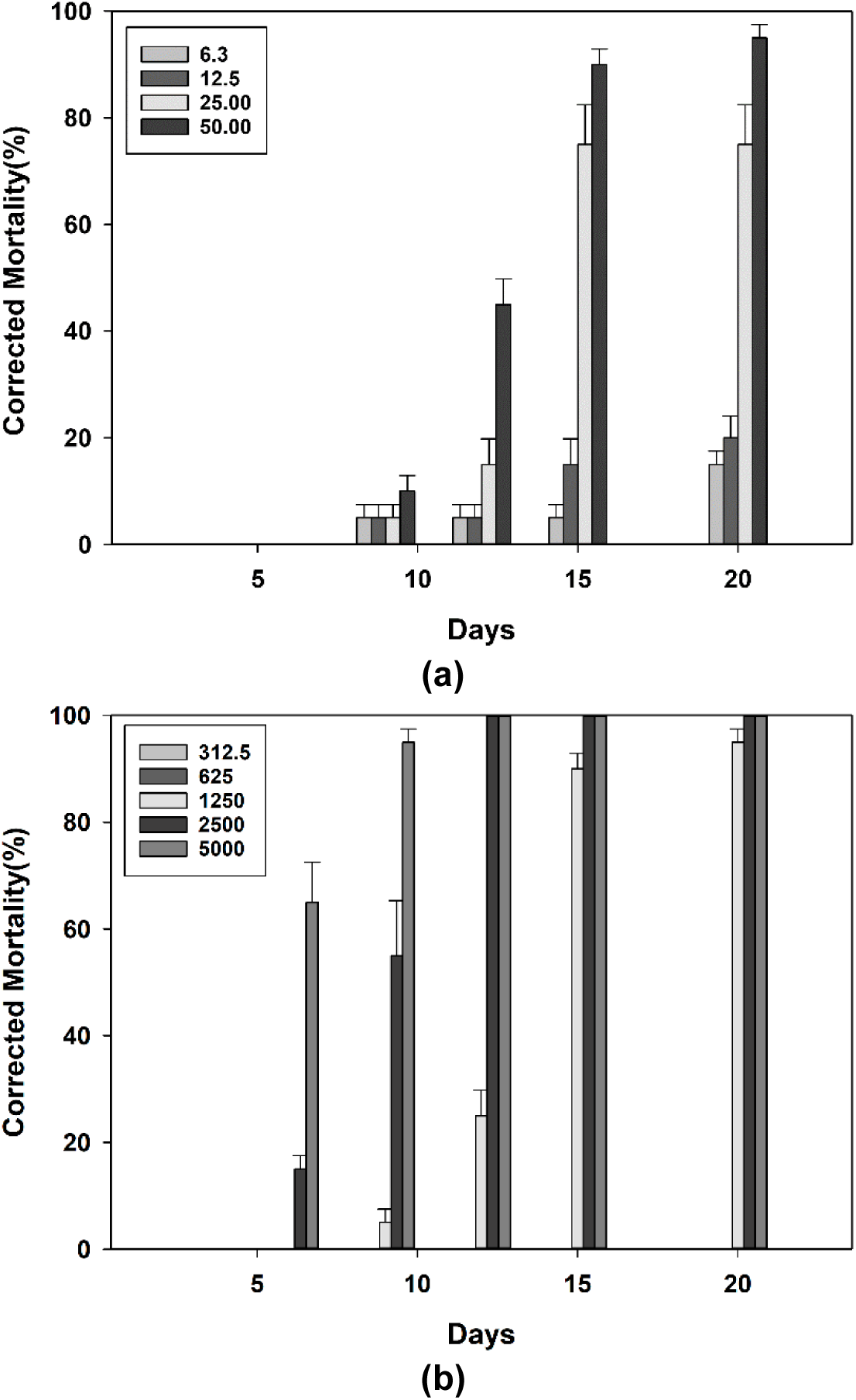
Effectiveness percentages (mean % ± SEM), as determined by the corrected mortality via Abbott’s formula, for *Rhynchophorus ferrugineus* larvae when fed a semi‒synthetic artificial diet containing: (a) neem oil or (b) gallotannin acid at different concentrations (ppm) under laboratory conditions (25 °C and 0:24 hours light/dark).

Adding neem oil to the semi‒diet of the *R. ferrugineus* larvae resulted in significant effects, as evidenced by the results of the omnibus test (chi-square likelihood ratio: 3000.208, df = 7, P < 0.001). Furthermore, both the dietary concentration factor (chi-square likelihood ratio: 70.288, df = 3, P < 0.001) and the covariate days (chi-square likelihood ratio: 975.549, df = 3, P < 0.001) presented significant statistical impacts. However, their interactions were not significant (chi-square likelihood ratio: 7.877, df = 3, P = 0.049). Neem oil caused 95.0±2.5% larval mortality effectiveness at 50.0 ppm by day 20, with lower dietary concentration being less effective (**Fig. 4 a**).

For the boric acid compound treatments of the *R. ferrugineus* larvae, significant effects were observed (omnibus test: chi-square likelihood ratio: 6408.077, df = 9, P < 0.001). The concentration factor (chi-square likelihood ratio: 4283.175, df = 2, P < 0.001), covariate days (chi-square likelihood ratio: 5788.338, df = 1, P < 0.001), and their interactions (chi-square likelihood ratio: 4346.817, df = 2, P < 0.001) also had significant impacts. Boric acid works faster and more efficiently. The corrected mortalities achieved with the two highest dietary concentrations (2500 and 5000 ppm) were 100% from day 12 of the trial (**Fig. 4b**).

Finally, for *R. ferrugineus* and the chemical compound gallotannin acid, no significant effects of concentration factor (chi-square likelihood ratio: 1.487, df = 1, P = 0.915), day (chi-square likelihood ratio: 0.497, df = 1, *P* = 0.481), or interaction (chi-square likelihood ratio: 2.466, df = 5, *P* = 0.782). Larval corrected mortality due to gallotannin acid was low at 5.0±2.5% for a concentration of 5000 ppm from day 6.

Boric acid was the only compound that resulted in notable mortality for the *P. interpunctella* larvae, as indicated in the previous section (**Fig. 2**). For this compound and this species, significant effects on corrected mortality were identified (Omnibus test: chi-square likelihood ratio: 4889.635, df = 9, *P* < 0.001). Concentration factor (chi-square likelihood ratio: 458.554, df = 2, *P* < 0.001), covariate day (chi-square likelihood ratio: 234.810, df = 1, P < 0.001), and their interactions (chi-square likelihood ratio: 157.969, df = 2, *P* < 0.001) had statistically significant effects. The highest effectiveness (100%) was observed for the 5000 ppm from 9 DAT (**Fig. 5**).

**Figure 5:**
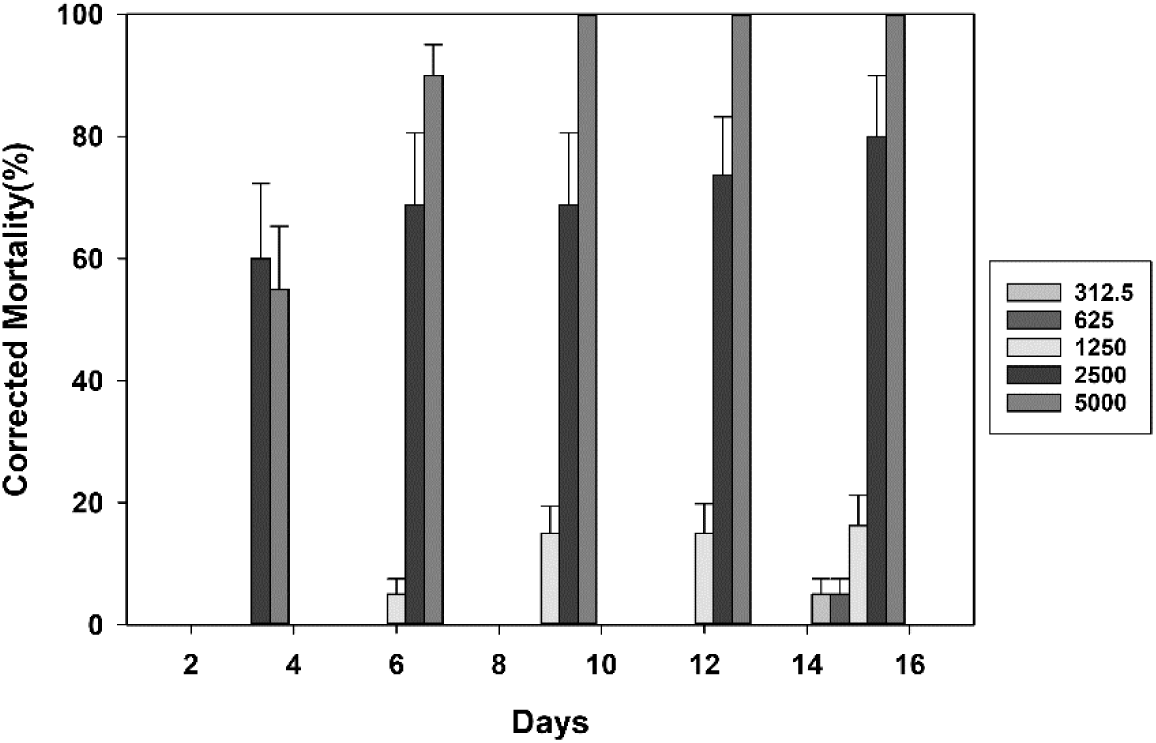
Effectiveness percentages (mean % ± SEM), as determined by the corrected mortality via Abbott’s formula, for *Plodia interpunctella* larvae when fed a semi-synthetic artificial diet containing boric acid at different concentrations (ppm) under laboratory conditions (25 °C and 0:24 hours light/dark).

**Fig. 6** presents the percentages of the *S. exigua* larval mortality effectiveness, which were calculated via Abbott’s formula [1], when the larvae were fed (a) neem oil, (b) boric acid, or (c) gallotannin acid.

**Figure 6:**
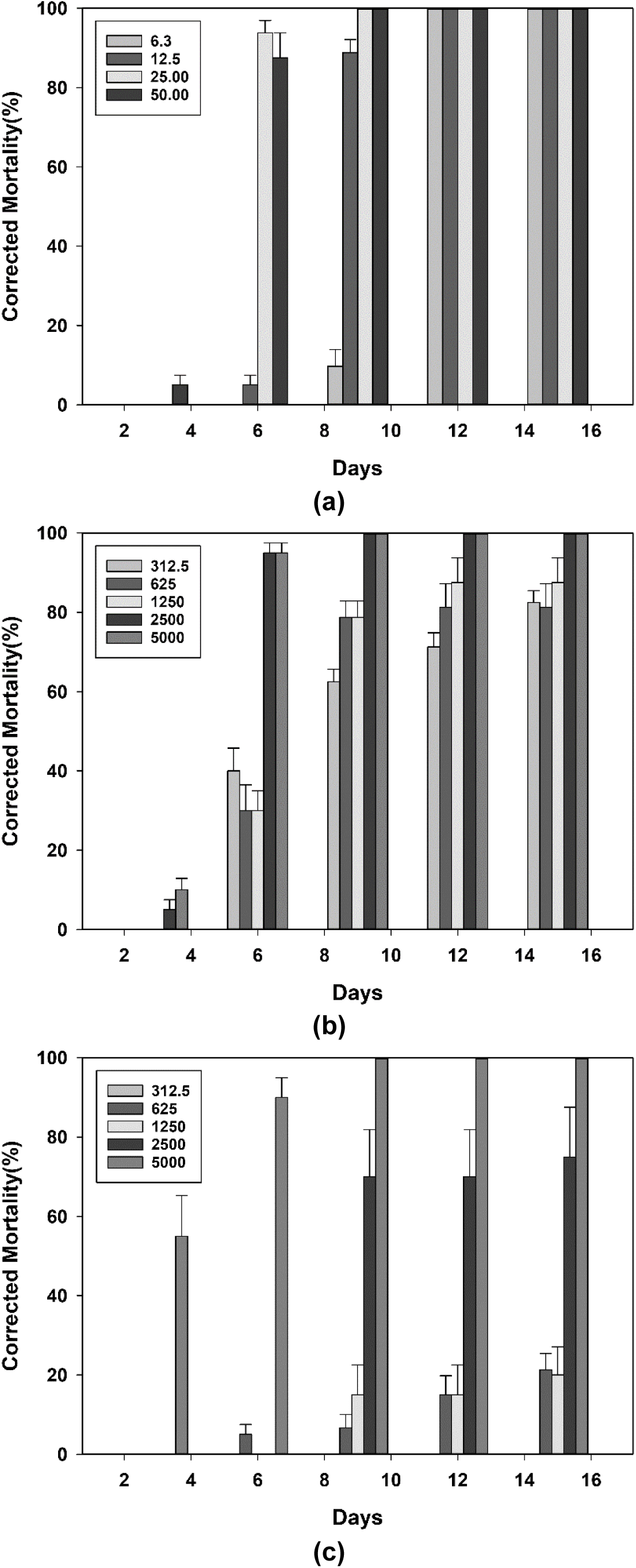
Effectiveness percentages (mean % ± SEM), as determined by the corrected mortality via Abbott’s formula, for *Spodoptera exigua* larvae when fed a semi-synthetic artificial diet containing: (a) neem oil, (b) boric acid, or (c) gallotannin acid at different concentrations (ppm) under laboratory conditions (25 °C and 0:24 hours light/dark).

Adding neem oil to the semi‒diet of the *S. exigua* larvae resulted in significant effects, as evidenced by the results of the omnibus test (chi-square likelihood ratio: 1876.828, df = 7, P < 0.001). Furthermore, the concentration factor (chi-square likelihood ratio: 316.720, df = 3, P < 0.001), the covariate days (chi-square likelihood ratio: 1697.510, df = 1, P < 0.001), and their interactions presented significant statistical impacts (chi-square likelihood ratio: 222.458, df = 3, P < 0.001). Neem oil caused complete larval mortality at 25 and 50 ppm by day 9, and at 6.3 and 12.5 ppm by day 12 (**Fig. 6a**).

The inclusion of boric acid in the semi‒diet led to significant effects, as evidenced by the results of the omnibus test (Chi-square likelihood ratio: 1679.304, df = 9, P < 0.001). The concentration factor (chi-square likelihood ratio: 214.121, df = 4, P < 0.001), day (chi-square likelihood ratio: 1410.769, df = 1, P < 0.001), and their interaction (chi-square likelihood ratio: 96.495, df = 4, P < 0.001) also presented significant impacts. This compound caused complete larval mortality at 2500 and 5000 ppm by day 9 (**Fig. 6b**).

Gallotannin acid treatment had significant effects (chi-square likelihood ratio: 6196.997, df = 11, P < 0.001). The concentration factor (chi-square likelihood ratio: 4322.744, df = 4, P < 0.001), day (chi-square likelihood ratio: 4075.252, df = 1, P < 0.001), and their interaction (chi-square likelihood ratio: 4019.120, df = 4, P < 0.001) also had significant impacts. This compound caused complete larval mortality at 5000 ppm by day 9 (**Fig. 6c**).

### 3.3. Summary of results

We can summarize the results found follows:

1. For the neem oil compound (azadirachtin), at the various concentrations used, the results revealed a high effectiveness (corrected mortality) rate for larvae of the species *S. exigua* (100% maximum mortality) for four concentrations). The effectiveness on the *R. ferrugineus* larvae was slightly lower (95.0±2.5% larval mortality effectiveness at 50.0 ppm on day 20). However, for the species *P. interpunctella*, no significant differences in mortality were detected.
2. Boric acid significantly affected the larval survival of all three species. The effectiveness values, measured by corrected mortality, were 100 percent in the three species from day 12 in the *R. ferrugineus* larvae and from day 9 for the *P. interpunctella* and *S. exigua* larvae.
3. For the gallotannin acid compounds, the results seem to present a different pattern to those of the other two chemical compounds. No significant effects on larval survival were found for this compound in *P. interpunctella* larvae, nor in effectiveness in the *R. ferrugineus* larvae. Only the *S. exigua* larvae seem to present a corrected mortality, with a pattern similar to that presented by the other two tested compounds. The maximum 100 percent was presented at 9 days and at a concentration of 5000 ppm.

## 4. DISCUSSION

At the larval level, neem oil is a chemical that has antifeeding effects (antixenosis) affecting insects through chemoreception (the primary effect) and by reducing their food consumption (the secondary effect). Moreover, it affects the synthesis of vital proteins in the digestive system of insect larvae. The increase in the concentration of this compound results in a higher degree of larval dysfunction and mortality (Dhra et al., 2018; Kilani-Morakchi et al., 2021). Commercial compounds containing this active ingredient (azadirachtin) are used to control various crop pest species, including Coleoptera, Hemiptera, Diptera, Orthoptera, and Isoptera (Morgan, 2009; Yasika & Shivakumar, 2025).

According to the results obtained for neem oil in our study, the concentrations used had greater effects on the *S. exigua* larvae (**Fig. 3a**) and slightly lower effectiveness on *R. ferrugineus* larvae (**Fig. 1a**). No response was observed in the *P. interpunctella* species. The neem oil values in *S. exigua* are similar to those reported by Medina et al. (2001) and Aggarwal et al. (2006).

A variety of insecticides have been the subject of experimental laboratory and field trials worldwide seeking to control *R. ferrugineus*. Of these, neem oil appears to show real potential for controlling this pest species (Naveed et al., 2023) and other species of the genus *Rhynchophorus* (Gabr et al., 2022) when used as active compounds. The corrected mortality results obtained for neem oil (**Fig. 4a**) were better than those reported in earlier works (Barranco et al., 1998, Merghem & Mohamed, 2017). The reason for this difference might be attributable to the use of different commercial products in the different bioassays.

Finally, with respect to neem oil, our results did not reveal significant effects on the mortality of *P. interpunctella* larvae caused by this compound which may be due to the concentration used. Nouri-Ganbalani et al. (2016) reported mortalities ranging from 20-80% but at relatively extremely high concentrations (from 100‒400 ppm). This is corroborated by the work of Rharrabe et al. (2008), who reported that at lower concentrations (2 ppm and 4 ppm), the mortality effects were mainly observed in pupae and adults of the species (25 and 34%, respectively).

With respect to boric acid, we detected effects of this compound on larval survival in three species: *R. ferrugineus* (**Fig. 1b**), *P. interpunctella* (**Fig. 2**), and *S. exigua* (**Fig. 3b**). This resulted in an effectiveness (corrected mortalities) of 100% at 9 or 12 days for all species at the highest concentration tested (5000 ppm): *R. ferrugineus* (**Fig. 4b**), *P. interpunctella* (**Fig. 5**), and *S. exigua* (**Fig. 6b**).

There are no bibliographic references on the mode of action and/or effectiveness of boric acid on the three species studied. However, according to previously published work, on *Blattella germanica* (L.) (Blatt.: Ectobiidae), was shown that boric acid’s mode action was to cause the insects to ultimately die of starvation. Examination of their intestines on days 1 and 2 post treatment revealed empty and slightly enlarged intestines. By days 3 and 4, the intestines were empty, enlarged, and filled with gas bubbles, with food intake under 1 mg per insect and a 50% mortality rate at day 3.8 (Cochran, 1995). Habes et al. (2006) studied the same species and reported that boric acid exhibited insecticidal effects. Their histological research suggested that death may have resulted from alterations in the midgut structure, and they confirmed neurotoxic impact of boric acid on this species. Similarly, in the ant species *Linepithema humile* (Hym.: Formicidae), electron microscopic studies revealed that ants fed low concentrations of boric acid (0.5%) presented gross abnormalities in the microvilli and cells lining the midgut (Klotz et al, 2002). More recently, in the larvae, pupae and adults of *Aedes aegypti* (Dip.: Culicidae), boric acid has been reported to cause malformations as well as mortality (Sharawi, 2023).

The above findings may explain our results; however, owing due to the biocidal nature of boric acid (McDonnell, 2017; Celikezen & Sahin, 2023), another hypothesis could be postulated that adds to boric acid’s mode of action.

From another perspective, certain specialized cells within insects, known as bacteriocytes, have endosymbiotic organisms or microbial structures. Others are ectosymbionts ‒ they are found on the body surface or on the surface of internal organs. Symbionts, which include mainly actinomycete fungi and bacteria, play a fundamental role in insect nutrition, as they allow many species to develop normally even though their food has limited nutritional value. Many symbiotic relationships are causal, especially for ectosymbionts, which often constitute rich gut microbiota (Thompson & Simpson, 2003). All insects (including insect pests) have symbiotic bacteria inside their body, particularly those insects that feed on restricted diets such as plant sap, vertebrate blood, or woody material. The symbionts play a prominent role in insect ecology. They aid in food digestion or provide nutrients; influence insect‒plant interactions, the host populations, heat tolerance, and pesticide detoxification; and as protect against natural enemies (Kashkouli et al., 2021). The syntrophic relationship within ectosymbiosis provides an important ecological evolutionary advantage (e.g., Gupta & Nair, 2020; Kashkouli et al., 2021; Drishnan et al., 2024). However, knowledge regarding various aspects of these associations is yet to be fully investigated (Kashkouli et al., 2021).

It has already been documented that palm weevils (including *R. ferrugineus*) host a diverse array of protozoan, fungal, viral, and bacterial species (Lefevre et al., 2004; Hoddle et al., 2024). This is because they have feed on materials, such as palm stipes, which has limited nutritional value (as mentioned before); thus, the ectosymbionts offer them a much-needed microbiota. The gut microbiome of *R. ferrugineus* comprises facultative and obligate anaerobic bacteria that are metabolized through fermentation. These bacteria are assumed to manage the palm tissue fermentation in tunnels where larvae develop and might play a key role in providing nutrition to insects (Tagliavia et al., 2014; Habineza et al., 2019). The contribution of microbial ectosymbionts to the physiology, reproduction, and detoxification of secondary plant metabolites in palm weevils, and in *R. ferrugineus* in particular, is not yet fully understood (Hoddle et al., 2024). Nevertheless, we do know how important this gut microbiota is in *R. ferrugineus* development (Butera et al., 2012; Habineza et al., 2019). The Lepidopteran species studied *P. interpunctella* (e.g., Montagna et al., 2016; Mereghetti et al., 2017a) and *S. exigua* (e.g., Gao et al., 2018; Eski et al., 2018), present and important and diverse gut microbiota. In general, the above has been cited for species of the Lepidopteran order (Mereghetti et al., 2017b; Zhang et al., 2022).

Lastly, we discuss the results obtained for gallotannin acid. Tannins are a group of structurally complex polyphenols, that include hydrolysable tannins, proanthocyanidins (syn. condensed tannins), and phlorotannins (Suvanto et al., 2017). Its name refers to proteins that bind and precipitate (Bule et al., 2020). Gallotannin acid (commonly named tannic acid) is the best-known hydrolysable tannin. The role of polyphenols is well understood in plant‒insect interactions and defenses, including their structure, induction, regulation, and anti-feeding and toxicity effects (Singh et al., 2021). In our study, the *S. exigua* larvae had low survival rates when exposed to gallotannin acid in the semi‒diet (**Fig. 3c**) at the highest concentration (5000 ppm), causing complete mortality (**Fig. 6c**). Schultz (1989) noted similar effects in other insect species. This author reviewed the cases and modes of action of tannins on insects and classified them into three groups: (1) those with negative effects (consumption, assimilation, growth and survival) (16 cases); (2) those with no effects (consumption, digestion, growth, survival, fecundity) (16 cases); and (3) those with positive effects (consumption, survival and growth) (11 cases). Therefore, our findings might lead us to consider that the species *S. exigua* falls into the first group above. In contrast, the species *R. ferrugineus* and *P. interpunctella* are would fall into the second group.

Possible explanations exist for the tannin-related classification of phytophagous insect species. Behmer (2008) postulates that when the food available to an insect varies both in its nutritional and allelochemical composition (e.g., tannins), foraging decisions probably represent a trade-off between the benefits gained from obtaining an optimal balance of nutrients and the costs of ingesting allelochemicals that may be detrimental. This seems to be the cases in the work on the African migratory locust *Locusta migratoria* L. (Orth.: Acrididae) (Raubenheimer, 1992; Simpson & Raubenheimer, 2000). When the locusts were given feed containing the optimal amount and ratio of protein to carbohydrates, they are immune to the effects of gallotannin acid even when it is present in concentrations of up to 10%. However, as the protein-to-carbohydrate ratios and the feed concentrations became increasingly suboptimal, the concentration of gallotannin acid needed to adversely affect insect performance decreased. Although our study looked at distinct species from a dietary point of view, the gallotannin acid concentration in the semi‒synthetic artificial diet was well below 10% (15.0% crude protein on a dry weight basis). Furthermore, as noted in the Materials & Methods section, the semi‒ synthetic artificial diet contained good levels of protein and carbohydrates. All the above could explain the differences found between the dose-response patterns for gallotannin acid in *R. ferrugineus* (**Fig. 1c**) and *S. exigua* (**Fig. 6c**), with those found for the other two compounds.

Our findings on the three natural compounds and the three species suggest new avenues for investigating these natural products as well as their potential as insect pest control agents that have reduced toxicological and environmental impacts. Further studies are required to develop and validate these research lines.

## 5. CONCLUSIONS

This study demonstrated the varying effectiveness of three natural compounds —neem oil, boric acid, and gallotannin acid— against the larval stages of three pest species: *Rhynchophorus ferrugineus*, *Plodia interpunctella*, and *Spodoptera exigua*.

1. Neem oil was highly effective at controlling *S. exigua*, achieving 100% mortality at the tested dietary concentrations. For *R. ferrugineus*, it presented slightly lower mortality rates. However, for the *P. interpunctella* larvae no significant effect was observed.
2. Boric acid exhibited universal effectiveness, with 100% corrected mortality rates for all three species at its highest concentration, underscoring its broad potential as a biocidal agent.
3. Gallotannin acid presented species-specific results, significantly impacting *S. exigua* survival but having negligible effects on *R. ferrugineus* or *P. interpunctella*.
4. These results underscore the need for further research into the interactions between these compounds and the gut microbiota of pest species, which could provide insights into optimizing their effectiveness. While the laboratory results are encouraging, additional field studies and long-term assessments are essential to validate these findings in practical agricultural settings.

## Author Contributions

Conceptualization, T.C.; methodology, T.C.; software, M.G.; validation, T.C. and M.G.; formal analysis, M.G.; investigation, T.C.; resources, T.C.; data curation, T.C. and M.G.; writing-original draft preparation, T.C.; writing-review and editing, M.G.; visualization, T.C. and M.G.; supervision, T.C.; project administration, T.C.; funding acquisition, T.C. All authors have read and agreed to the published version of the manuscript.

## Data Availability Statement

The data that supports the findings of this study are available from the corresponding author, [TC] and the second author [MG], upon reasonable request.

